# Meta-analysis of behavioural research in lizards reveals that phylogeny and viviparity contribute better to animal personality than secretory glands

**DOI:** 10.1101/2023.05.12.540450

**Authors:** M.R. Ruiz-Monachesi, J.J. Martínez

## Abstract

Animal personality is defined as an individual’s behavioural consistency across contexts, situations, and time. Understanding the evolution of animal personality requires the integration of macroevolutionary patterns with intraspecific promoters of individual behavioural consistency. In this study, we conducted a meta-analysis to assess the association between lizards’ animal personality and different indicators of sociability (a personality promotor) in a phylogenetic context. In lizards, the presence of both, secretory glands and viviparity have been associated with higher sociability levels. We analysed behavioural repeatability data, including 490 effect sizes from 37 species and 63 studies, considering five categories (activity, aggressivity, boldness, exploration, sociability) while controlling for phylogenetic constraints. For each species, we obtained data on the number of secretory glands and the reproductive mode (oviparous or viviparous). The results showed similar values of repeatability for species with and without glands and an absence of correlation between the number of glands and repeatability data. However, higher repeatability was present in viviparous species than in oviparous species. When we conducted separate analyses for each behavioural type, we found two contrasting patterns for exploration and boldness. Species without glands were more exploratory, while species with glands were bolder. In general, phylogeny explained the observed patterns of repeatability, but boldness, exploration and sociability were poorly explained by evolutionary history among species. This study represents a first step in disentangling the integration among animal personality, life-history and morphology traits under a broad evolutionary context.

## 1-INTRODUCTION

Individual behavioural consistency, or animal personality, is a phenomenon where individuals within a population exhibit different and consistent behaviours over time and across contexts (Dall et al., 2004; Dingemanse et al., 2010; Réale & Dingemanse, 2012; Laskowski et al., 2022). From a statistical perspective, the behavioural variance within a population can be divided into two components: the among-individual and within-individual variance components. The former describes how different individuals are from each other, while the latter describes how consistent individuals are in repeated expressions of behaviour (Nakagawa & Schielzeth, 2010; Brommer, 2013). The extent of individual differences in behavioural or physiological scores maintained over time is called repeatability (Dochtermann & Dingemanse, 2013; Biro & Stamps, 2015), which is usually quantified using the repeatability (*R*) index (Nakagawa & Schielzeth, 2010). This index represents the amount of the total phenotypic variance that is due to differences between individuals measured considering different predictors (i.e., variation in time and external stimuli) and factors (e.g., sex, maturity; Biro & Stamps, 2015). The generation and maintenance of this variation have been broadly studied at the microevolutionary scale (e.g., Santostefano et al., 2021) being subject to natural selection and evolution (Dingemanse & Réale, 2005; Smith & Blumstein, 2008). However, macroevolutionary patterns and general trends can be observed considering a broad phylogenetic context (Hansen & Martins, 1996) and may allow for revealing different aspects of animal personality.

Macroevolutionary studies of animal personality are challenging to perform, particularly when considering a broad taxonomic sample. Data acquisition for this approach takes a long time, as it involves collecting data from different individual repetitions in natural or/and experimental conditions (Dingemanse & Wright, 2020). Furthermore, behaviours are more difficult to standardize across studies, which can limit the number of taxa studied. One possible approach to addressing macroevolutionary patterns and trends in animal personality is through meta-analytic studies. These allow for a broad taxonomic sample and help to infer evolutionary patterns (e.g., Bell et al., 2009; Garamszegi et al., 2012). Recently, different meta-analytic studies have shown the presence of animal personality on different behaviours, including cognitive abilities, individuals’ survival, spatial movements, and social learning (Cauchoix et al., 2018; Dougherty & Guillette, 2018; Moiron et al., 2020; Stuber et al., 2022; Camacho-Alpízar & Guillette, 2023), in addition to its heritability patterns (Dochtermann et al., 2019). Furthermore, others meta-analyses studies have evidenced the association of animal personality with different intrinsic and extrinsic factors (Laskowski et al., 2022). For instance, intrinsic factors that modify an individual’s behavioural responses (e.g., metabolism, hormones, energetic reserves) appear to present weak associations with animal personality (Niemelä & Dingemanse, 2018), although differences in growth rates between individuals may favour differences in their personalities (Stamps, 2007). Animals that are relatively aggressive, bold, explorative and/or active may also have proportionally high metabolic rates, hormone levels, body weights, and/or body sizes (Niemelä & Dingemanse, 2018). Extrinsic factors, such as behavioural types, taxonomic groups (i.e., invertebrates vs. vertebrates), the conditions where data were obtained (field/laboratory), the interval between observations (short vs. long), age (juveniles vs. adults) and sex (but see Szabo et al., 2019; Harrison et al., 2022) could also explain the differences in the individual consistency among taxa (Bell et al., 2009). Predators may also influence an individual’s consistency, particularly in bolder individuals, favouring their learning processes (Dougherty & Guillette, 2018).

The mechanisms involved in the evolution of animal personality arise from conspecifics interactions, such as sexual interactions (Dingemanse & Réale, 2005; Réale & Dingemanse, 2010; Schuett et al., 2010) or non-sexual interactions (i.e., interactions without reproductive intentions; Gartland et al., 2022). Among non-sexual interactions, social relationships are particularly relevant, and the social niche specialization hypothesis proposes that social conflicts and alternative social options play an important role in the evolution of animal personality (Bergmüller & Taborsky, 2010; Montiglio et al., 2013). In this sense, more social animals may also exhibit higher individual behavioural consistency. For example, more social shrew species demonstrate higher individual consistency (aggressiveness) in social contexts compared to less social species (von Merten et al., 2017). Similarly, crickets, exposed to conspecifics exhibit higher individual consistency than solitary ones (Jäger et al., 2019).

Lizards manifest different degrees of sociability and animal personality, as was evidenced by revisions such as Gardner et al. (2016) and Waters et al. (2017). Although meta-analytic approaches have been employed in lizards’ studies (e.g., Ashton & Feldman, 2003; Luiselli, 2008; Noble et al., 2018; Szabo et al., 2019; Doherty et al., 2020; Putman & Tippie, 2020; Tan et al., 2023), there are limited studies on animal personality in this group. One relevant study by Putman and Tippie (2020) found that urbanization in lizards did not have an impact on lizards’ behavioural types (activity, exploration, boldness, sociability). Moreover, there are no meta-analytic studies evaluating animal personality under the social niche specialization hypothesis in lizards. A recent study by Baeckens and Whiting (2021) explored the relationship between sociability and the evolution of chemical secretory glands in lizards, suggesting that the investment in these glands may indicate sociability and facilitate its evolution. Viviparous lizard species have been shown to frequently exhibit stable social aggregations (Gardner et al., 2016) and live-bearing may promote the evolution of sociability in these taxa (Halliwell et al., 2017) due to the formation of permanent social groups or groups in which individuals maintain permanency across multiple seasons.

Based on the social niche specialization hypothesis, we conducted a meta-analytic study on repeatability data in lizards to investigate the relationship between two putative sociability indicators: the number (or presence) of chemical secretory glands and the reproductive mode on individual behavioural consistency in lizards. Firstly, we expect that species with a greater number of secretory glands (or the presence of these) will show higher individual consistency than those with fewer glands (or the absence of these). Secondly, we predict that viviparous species will exhibit higher individual consistency than oviparous species.

Furthermore, various factors can affect animal personality (e.g., Bell et al., 2009; Schuett et al., 2010; Niemelä & Dingemanse, 2018). However, the effects of these factors on animal personality depend on the behavioural type analysed and the phylogeny (e.g., Garamszegi et al., 2012; Ducatez et al., 2017; Putman & Tippie, 2020; Queller et al., 2022). Therefore, we conducted conventional and phylogenetic meta-analyses on separated behavioural types and pooled data to examine the influence of phylogeny on the repeatability of different behavioural types (Réale et al., 2007).

## 2 MATERIAL AND METHODS

### 2.1 Literature search

We conducted a comprehensive literature review to perform a meta-analysis on lizards’ behavioural consistency, using the *R* index as our primary measure. We searched in Google Scholar, Elsevier and ScienceDirect databases for studies published up to March 2023, using the search string: “behavioural types* OR intraclass coefficient* OR repeatability* OR personality AND lizards*”. On these studies, we scanned all titles with lizards’ data for relevance, focusing on personality or repeatability/intraclass coefficient data. A total of 80 relevant studies were identified, comprising 76 published articles and 4 theses. From these, we found repeatability data in 62 articles and calculated 6 repeatability values using the “*rpt*” function from the package *rptR* (Stoffel et al., 2017) for one study (Carazo et al., 2014). Therefore, our final dataset consisted of 490 repeatability values from 62 articles published in 30 different journals and one thesis.

### 2.2 Data preparation

We extracted data on repeatability, or the intraclass coefficient of correlation, and classified them into five behavioural types: *i*- activity, *ii*- aggressiveness, *iii*-boldness, *iv*- exploration, and *v*-sociability, following Réale et al.’s (2007) classification. We incorporated information about different factorial moderators (“Age”, “Behavioural type”, “Data Collected”, “Sex”, “Stimuli”, “Time” and “Treatment Binary”; Table 1) and numerical moderators (“Mean of repetition per individual” and “Sample Size”; Table 1) that we expected *a priori* being influent on responses variable based on our reading of these studies.

**Table 1:** List and definition of the different moderators obtained from the literature revision, considering those factorial (“Age”, “Behavioural type”, “Data collected”, “Sex”, “Stimuli”, “Time” and “Treatment Binary”) and numerical (“Mean number of assays per individual”, “Sample size”) moderators.

| Moderator | Definition | Levels |
| --- | --- | --- |
| Age | Individuals ontogenetic stage. | Adults -Juveniles |
| Behavioural type | Behavioural types based on Réale et al. (2007). | Activity -Aggressiveness-Boldness-Exploration-Sociability |
| Data Collected | Conditions of data obtention. | Field-Laboratory- Semi-natural |
| Mean reps. per ind. | Mean number of assays per individual. |  |
| Sample Size | Sample size considered in the study. |  |
| Sex | Sex of individuals. | Males-Females |
| Stimuli | Main stimulus used to trigger the behavioural response. | Food, Non-novelty, Novelty, Predatory,<br>Risk-taking, Social- Thermic |
| Time | Period of time of the study. | Days-Weeks-Months- Years |
| Treatment Binary | i.e., whether the data provide treatment or control conditions. | Control-Treatment |

For each species with repeatability data on personality traits, we obtained the average number of chemical secretory glands (N° of glands) and the reproductive mode of the species (i.e., oviparous or viviparous) from various sources (see Supplementary Table S1).

### 2.3 Phylogenetic tree

As different lizard species share a common phylogenetic history, they may not be considered independent statistical units (Felsenstein, 1985). Therefore, we made phylogenetic corrections using the Squamate’s molecular phylogeny from Zheng and Wiens (2016). As not all studied species were present in this phylogeny, we performed a metatree using Mesquite v2.74 (Maddison & Maddison, 2017) and included three missing species studied herein (*Egernia striolata*, *Lampropholis similis*, *Podarcis virescens*). We set branch lengths equal to one and subsequently fitted the tree to ultrametricity using the “nnls.tree” function of the *phargon* package (Schliep et al., 2021). From this tree (Figure 1) we obtained a correlation squared-matrix using the “vcv” function from the *ape* package (corr=TRUE; Paradis et al., 2004). We used this matrix (i.e., phy cor-matrix) as a random effect in the subsequent analysis to control for the shared evolutionary history of studied species (Nakagawa & Santos, 2012).

**Figure 1:**
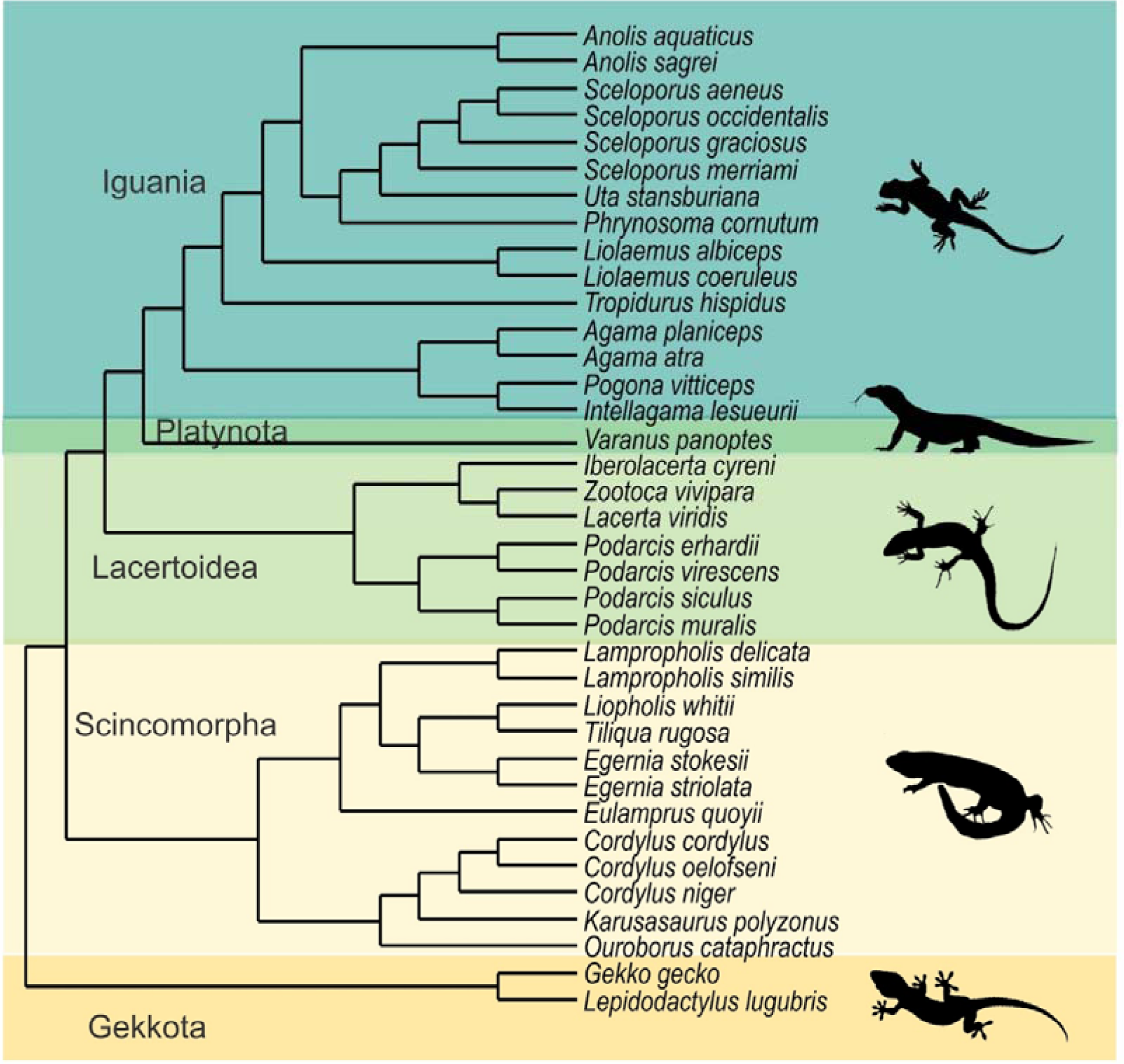
Lizard phylogenetic metatree based on molecular analysis by Zheng and Wiens (2016), showing the phylogenetic relationships among 37 studied species.

### 2.4 Meta-analysis

Following Stuber et al. (2022) and Holtmann et al. (2017), we calculated the effect size of repeatability data (henceforth called “*Zr*”) by performing Fisher’s Z-transformed standardization. We used *Metafor* package v. 3.8.1 (Viechtbauer, 2010) for all the following analyses.

First, we ran a null-random model (Borenstein et al., 2010; Riley et al., 2011), i.e., without specifying random variables, employing the “rma.uni” function (method = "REML") on the *Zr* variable. This approach allowed testing for heterogeneity (*Q*; Cochran, 1954) present in *Zr*, where a significant *Q*-value suggests the presence of more variability than expected by sampling bias. Additionally, from this null-random model, we obtained the amount of heterogeneity (*τ*^2^) and reported the percentage of heterogeneity (*I^2^*; Higgins & Thompson, 2002; Viechtbauer, 2005). On this null-random model, we performed Egger’s test (to check for funnel plot asymmetry; Sterne et al., 2005) and Rosenthal’s failsafe test to detect for publication bias (Becker, 2005). Additionally, we made a histogram to observe the frequency of data.

Second, we used the function “rma.mv” and ran a multivariate random-effects model (Borenstein et al., 2010), specifying “study identity” (Study), “the number of observations” (obs) and “the phylogeny” (species.id.phy) as random variables. The model was: *Zr*, random = (∼ 1 | Study/obs, ∼ 1 | species.id.phy), R=list (species.id.phy=phy cor-matrix). This approach allowed us to estimate the variance and percentage of contribution from each variable to the total random variability.

Third, we ran a mixed effects model test, exploring factorial and numerical moderators (Table 1) and random effects that may affect our data. From this model, we obtained a square variance matrix used in the following analyses to control these moderator effects (called by us fixed controlled variability) used for control.

Fourth, we used “Occurrence of chemical glands” (i.e., absence/presence), “Number of chemical glands” and “Reproductive mode” (oviparous/viviparous) as moderator variables and ran three models, each including one of these three as an independent variable. While the random variables were: Study*obs, phylogeny, and the fixed controlled variability.

Additionally, we explored whether repeatability and moderators (occurrence of chemical glands, number of chemical glands, and reproductive mode) present different patterns considering each behavioural type separately (activity, aggressiveness, boldness, exploration, sociability; Supplementary S2).

Finally, for all mentioned models, we ran the same models without adding phylogenetic information as a random effect to observe the effect of phylogeny on data. In all models, we obtained the back repeatability (*R*) and its confidence intervals (95% CI) following Stuber et al. (2022).

All mentioned analyses were performed using R v. 4.2.3 (R Core Team, 2023).

## 3 RESULTS

### 3.1 Descriptive analysis of compiled research

We obtained data for 37 species within 24 lizard genera (Figure 1), representing five superfamilies: Gekkota (N_study_ = 3; 4.84%), Iguania (N_study_ = 18; 29.03%), Lacertoidea (N_study_ =15; 24.19%), Platynota (N_study_ = 1; 1.61%) and Scincomorpha (N_study_ = 25; 40.33%). The families most studied were Scincidae and Lacertidae (N_study_ =23; 37.10%, N_study_ = 15; 24.19 %; respectively) and the lesser ones were Cordylidae, Gekkonidae, Liolaemidae and Varanidae, each one with one, two or three studies. The most studied genus was *Lampropholis* (N_study_ =12; 19.05%) followed by *Zootoca*, *Iberolacerta*, *Liopholis*, *Podarcis* and *Tiliqua*, each one with four/five studies. *Lampropholis delicata* was the most extensively studied species, with ten studies, followed by *Zootoca vivipara* with five studies. The two genera with the most repeatability data were *Lampropholis* (N_data_ =122; 24.90%) and *Podarcis* (N_data_=68; 13.88%). *Iberolacerta*, *Sceloporus* and *Zootoca* presented between 30-31 repeatability values each one.

Boldness (N_data_ = 200; 40.82%) and exploration (N_data_ = 151; 30.82%) constituted the majority of repeatability data. Aggressiveness (N_data_ = 57; 11.63%), activity (N_data_ = 48; 9.8%) and sociability (N_data_ = 34; 6.94%) contributed less to the totality of data.

### 3.2 Null random model

A total of 490 repeatability values were included in this analysis, and observed outcomes ranged from 0.0 to 2.242. The model was significant (F_df1, 489_ = 964.47, p < 0.001) and evidenced a high heterogeneity across data (*Q* _df489_ = 3639.691, p < 0.001; τ^2^ = 0.107; *I* ^2^= 86.39%; Table 2), suggesting more variability present than those expected by sampling bias (*H*^2^ = 7.35). The average of *Zr* was 0.536 [95% CI 0.502-0.571] and differed significantly from zero (z = 31.056, p < 0.001).

**Table 2:** Summary showing the number of: studies (N_Study_), effect size (N_Effect size_) and species (N_Sp._) considered per behavioural type (i.e., Activity, Aggressiveness, Boldness, Exploration and Sociability) and for the pooled dataset. Parameters of null variables random model: *Q* = Cochran ’s Q or heterogeneity value; *τ*^2^ = amount of heterogeneity of the model; *I*^2^ = percentage of variation across studies that is due to heterogeneity; *H*^2^= ratio of variability due to of heterogeneity over the variability due to sampling bias, higher than one values suggest fewer sampling bias. Variance of the total random model (*σ*^2^_Total random_) and the percentage of random variability accounted by phylogeny (*σ*^2^_Total random_), studies (*σ*^2^_Total random_), interaction of study/observation (*σ*^2^_Total random_) and fixed controlled variables (*σ*^2^_Total random_). Additionally, it shows the effect size and repeatability values with 95% confidence intervals. * Did not incorporate in this analyse.

| Data | Activity | Aggressiveness | Boldness | Exploration | Sociability | Pooled |
| --- | --- | --- | --- | --- | --- | --- |
| $N_{\text{Study}}$ | 10 | 16 | 45 | 26 | 12 | 62 |
| $N_{\text{Effect size}}$ | 48 | 57 | 200 | 151 | 34 | 490 |
| $N_{\text{Sp.}}$ | 7 | 10 | 34 | 17 | 6 | 37 |
| $Q$ | 645.70 | 316.740 | 809.080 | 642.464 | 460.16 | 3639.691 |
| $\tau^2$ | 0.150 | 0.0721 | 0.109 | 0.080 | 0.076 | 0.107 |
| $I^2$ | 92.80% | 84.59% | 82.54% | 76.68% | 91.20% | 86.39% |
| $H^2$ | 13.88 | 6.49 | 5.73 | 4.29 | 11.37 | 7.35 |
| $\sigma^2_{\text{Total random}}$ | 0.254 | 0.131 | 0.120 | 0.081 | 0.088 | 0.301 |
| $\sigma^2_{\text{Phylogeny}}$ | 67.91% | 62.37% | 7.58% | 9.14% | 0% | 78.28% |
| $\sigma^2_{\text{Study}}$ | 0 | 10.71% | 0% | 0% | 33.75% | 0% |
| $\sigma^2_{\text{Study/obs}}$ | 33.08% | 5.80% | 5.50% | 74.20% | 66.25% | 21.72% |
| $\sigma^2_{\text{Fix contr. Var}}$ | 0 | 11.45% | 86.92% | 16.66% | 0% | * |
| Effect size | 0.528 [-0.018-1.076] | 0.664[0.316-1.012] | 0.568 [0.448-0.689] | 0.455[0.337-0.573] | 0.330[0.133-0.527] | 0.546 [0.114-0.974] |
| Repeatability | 0.484[-0.019-0.792] | 0.581[0.306-0.767] | 0.514[0.420-0.597] | 0.426[0.325-0.518] | 0.319[0.133-0.483] | 0.497[0.113-0.752] |

The data showed significant asymmetry according to Egger’s test (estimate = 0.675 [95% CI 0.62-0.73], t = −3.582, p < 0.01), as can be seen in the funnel plot (Figure 2A). The random effects model imputed 89 missing studies on the right side, resulting in an average *Zr* of 0.633 [95% CI 0.598-0.667], which is ∼ 1.18 higher than the obtained *Zr* of 0.536. This can be seen in Figure 2B, which shows higher frequencies of *Zr* in the first two quartiles and lower frequencies in the 2nd and 3rd quartiles (Figure 2B). Rosenthal’s failsafe N was 1,114,233, suggesting a greater confidence in the meta-analytic findings. This is the putative number of studies needs to avoid a bias in literature sampling. The overall repeatability for the null random model was *R*= 0.490 [95% CI 0.464-0.515].

**Figure 2:**
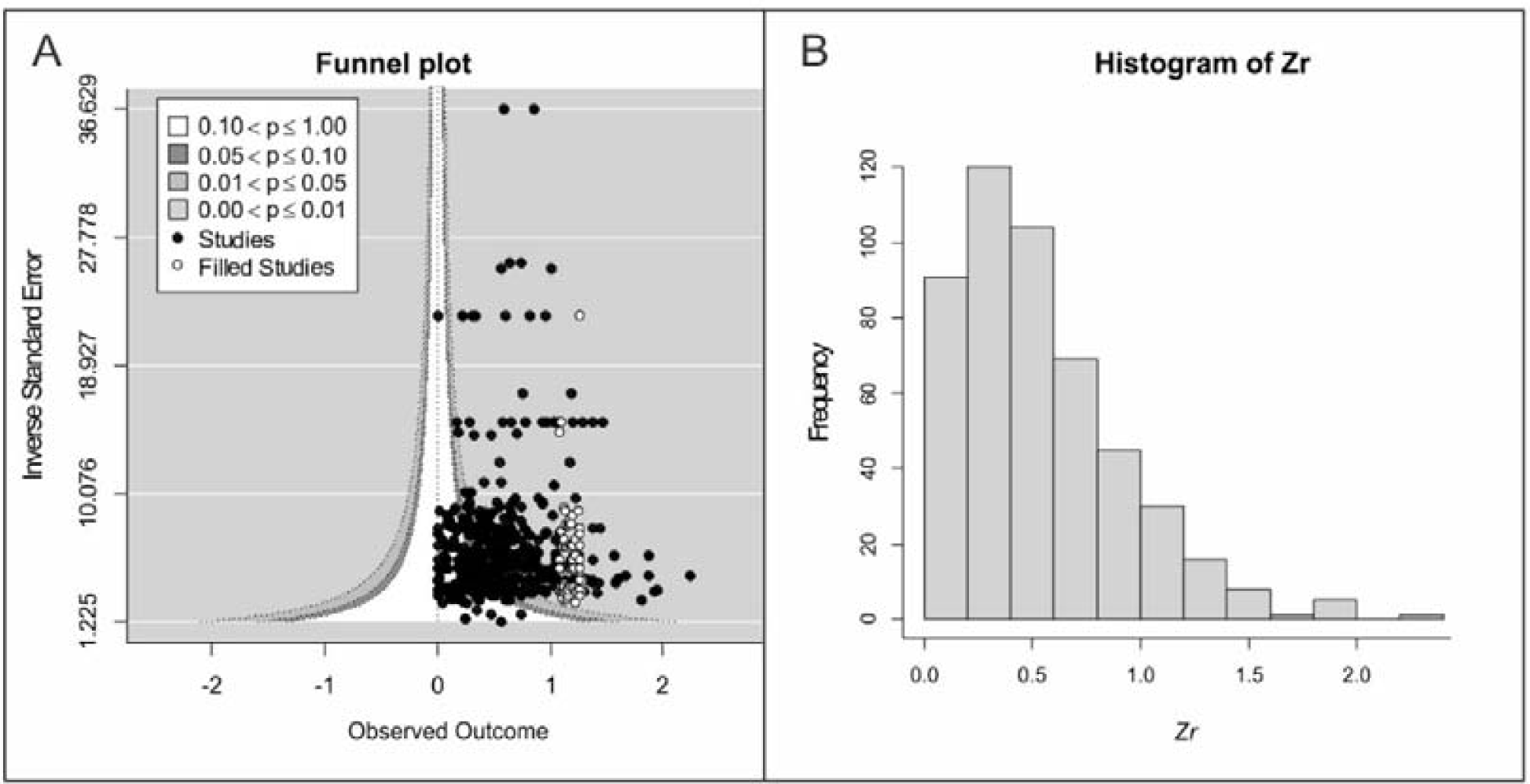
A) Funnel plot with right-hand side asymmetry data, white dots represent those potentials studies with an effect size around 1.5 and 2.2. B) Histogram draw of *Zr* values.

### 3.3 Random-effects model (summary in table 2)

The random variables model was found to be statistically significant (F _(df1= 1, df2= 36)_ = 6.56, p = 0.014). The total random variability (σ^2^_total random_) was 0.301, with phylogenetic relationships among species explaining 78.28% of random variability (σ^2^_phy_= 0.235) indicating a strong phylogenetic pattern on effect size data (Table 2, Figure 3). The study/observation factor accounted for 21.72% of random variability (σ^2^_study/obs_ = 0.065) indicating variation in repeatability among studies and observations. The average value of *Zr* was statistically significant (p = 0.014; *Zr =* 0.546 [95% CI 0.114-0.978]), and the repeatability for the random variable model was *R* = 0.497 [95% CI 0.113-0.752] as is shown in the Figure 3. A summary of these results can be observed in Table 2.

**Figure 3:**
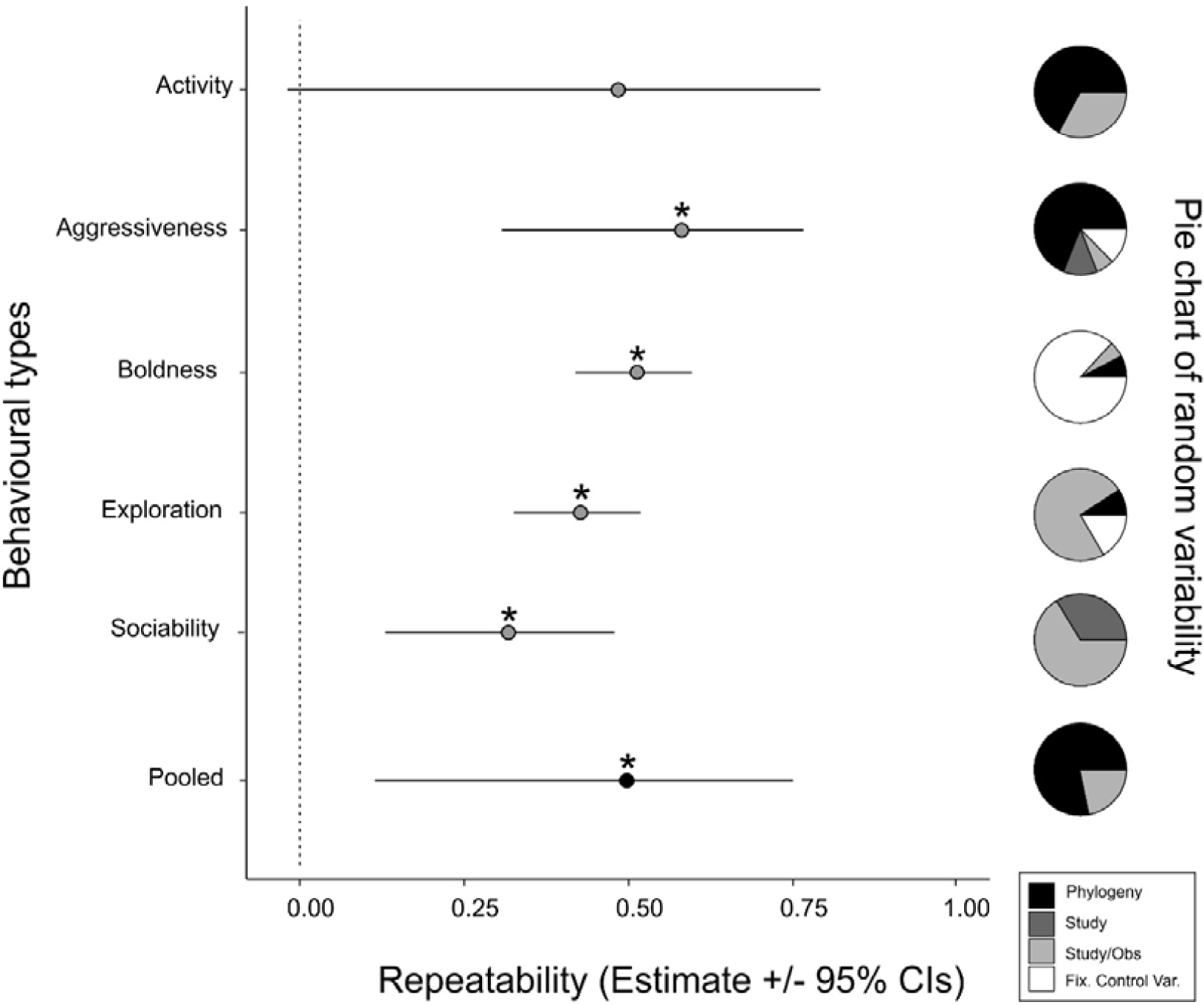
Repeatability values (± 95% CI) obtained for five behavioural types and pooled data. The asterisk indicates values statistically significant. The right panel shows pie charts with the percentage of random variability (Phylogeny, Study, Study/Obs, and Fixes controlled variables) which explain each behavioural type and pooled data.

### 3.4 Mixed effects model test

The mixed effects model was found to be statistically significant (F _(df1= 22, df2= 468)_ = 7.6026, p < 0.001). The total random variability (σ^2^_total random_) was 0.153, with 67.25% of the variability attributed to phylogeny (σ^2^_phy_= 0.103) and 32.42% attributed to observations (σ^2^_obs_ = 0.050). The fixed variables that were found to be significant were “Age”, “Mean Reps per Ind”, “Time” and “Treatment_binary”, and further details can be found in Tables 1 and 3. Comparison of these variables within their levels showed significant differences (Table 3).

**Table 3:** Fixed factors from full models (i.e., those models with all fixed and random variables incorporates), showing their: average effect size of repeatability (*Zr*), 95% CI low and upper boundaries (Lb, Ub, respectively) of *Zr* and p-values from a comparison of fixed variables levels factors. In bold, statistically significant values.

| **Moderators** | ***Zr*** | **Lb** | **Ub** | ***p*** |
| --- | --- | --- | --- | --- |
| **Factor (Age) Adults** | **0.483** | **0.036** | **0.929** | **0.034** |
| **Factor (Age) Juveniles** | **0.486** | **0.018** | **0.953** | **0.042** |
| Scale (Sample_Size) | -0.024 | -0.058 | 0.009 | 0.155 |
| **Scale (mean_reps_per_ind)** | **0.185** | **0.145** | **0.225** | **<0.001** |
| Factor (Data_Collected) Lab | -0.050 | -0.226 | 0.126 | 0.574 |
| Factor (Data_Collected) Semi-nat | 0.052 | -0.216 | 0.319 | 0.704 |
| **Factor (Time) Months** | -0.029 | -0.132 | 0.074 | 0.577 |
| Factor (Time) Weeks | 0.010 | -0.096 | 0.115 | 0.857 |
| **Factor (Time) Years** | **0.246** | **0.072** | **0.420** | **0.006** |
| **Factor (Treatment_binary) T** | **0.102** | **0.015** | **0.188** | **0.021** |
| Factor (Sex) M | -0.027 | -0.143 | 0.089 | 0.649 |
| Factor (Sex) M-F | -0.098 | -0.224 | 0.027 | 0.124 |
| Factor (Behavior_type) Aggressiveness | 0.166 | -0.116 | 0.448 | 0.247 |
| Factor (Behavior_type) Boldness | -0.024 | -0.260 | 0.212 | 0.840 |
| Factor (Behavior_type) Exploration | -0.053 | -0.295 | 0.189 | 0.667 |
| Factor (Behavior_type) Sociability | -0.227 | -0.542 | 0.089 | 0.159 |
| Factor (Stimuli) Food | -0.134 | -0.424 | 0.156 | 0.365 |
| Factor (Stimuli) Non-noventy | 0.121 | -0.183 | 0.424 | 0.435 |
| Factor (Stimuli) Noventy | 0.139 | -0.078 | 0.357 | 0.209 |
| Factor (Stimuli) Predatory | -0.007 | -0.194 | 0.180 | 0.942 |
| Factor (Stimuli) Risk-taking | -0.023 | -0.228 | 0.182 | 0.827 |
| Factor (Stimuli) Thermic | 0.127 | -0.114 | 0.368 | 0.301 |

### 3.5 Separated behavioural types

Details in the supplementary data S1 and Table 2; boldness evidenced the highest heterogeneity (*Q* = 809.080), while aggressivity had the lowest one (*Q* = 316.740; Table 2). Activity and sociability showed a higher percentage of variation across studies (*I^2^* = 92.80% and 91.20%, respectively) and exploration was the lowest (*I^2^* = 76.68%; Table 2). All five behavioural types showed values of *H*^2^ that deviated significantly from one (Table 2), suggesting that variability across studies is not due to a sampling bias. The phylogeny explained strongly the observed pattern in activity and aggressiveness (more than 60 % in both, Table 2, Figure 3). In contrast, phylogenetic relationships explained less variation for boldness and exploration and did not explain sociability (Table 2, Figure 3). Variability in exploration and sociability was largely explained by differences across studies and observations (Table 2, Figure 3), while fixed controlled variables highly explained boldness (86.92%, Table 2, Figure 3) whereas aggressiveness and exploration were subtly explained by it (11.41% and 16.66%, respectively; Figure 3).

All repeatability values were significant, except by the activity (*R =* 0.484 [95% CI - 0.019-0.792]; Table 2; Figure 3). Higher repeatability values were for aggressiveness and boldness (*R* = 0.581 [95% CI 0.306-0.767]; *R* = 0.514 [95% CI 0.420-0.597], respectively; Table 2; Figure 3), while lower values were for sociability and exploration (*R =*0.319 [95% CI 0.133-0.483], *R =*0.426 [95% CI 0.325-0.518], respectively; Figure 3).

### 3.6 Glands and viviparity

The model considering the occurrence of chemical glands (OCG) was marginally not significant (F _(df1= 2, df2= 35)_ = 3.243, p = 0.051). The σ^2^_total random_ was 0.306, with phylogeny accounting for 77.78% of the variability (σ^2^_phy_ = 0.238), observations accounting for 19.31% of the variability (σ^2^_obs_ = 0.059), and fixed controlled variability accounting for 2.78% (σ^2^_fix. contr. var_ = 0.009). The average *Zr* for “Glands Absent” was marginally not significant (p =0.054; *Zr* = 0.530 [95% CI −0.009-1.067], Table 4), whereas for “Glands Present” it was statistically significant (p=0.02; *Zr* = 0.554 [95% CI 0.096-0.985]; Table 4). Posterior comparison found no significant differences between the two groups (t= 0.107, p = 0.91). The overall repeatability for “Glands Absent” group was *R*_GAbs_= 0.485 [95% CI −0.008-0.788] (Figure 4; Table 5), while that of the “Glands Present” group was *R*_GPres_= 0.503 [95% CI 0.096-0.766] (Figure 4; Table 5). The model excluding the phylogenetic effect found significant *Zr* for both “Glands Absent” (p < 0.001; *Zr* = 0.533 [95% CI 0.457-0.609]; Table 4) and “Glands Present” (p < 0.001; *Zr* = 0.561 [95% CI 0.496-0.625]; Table 4) groups. Posterior comparison found no significant differences between both groups (t = 0.54, p = 0.58). The overall repeatability of the “Glands Absent” group was *R*_GAbs_= 0.488 [95% CI 0.428-0.543] (Figure 4; Table 5), while that of the “Glands Present” group was *R*_GPres_= 0.508 [95% CI 0.459-0.555] (Figure 4; Table 5).

**Figure 4:**
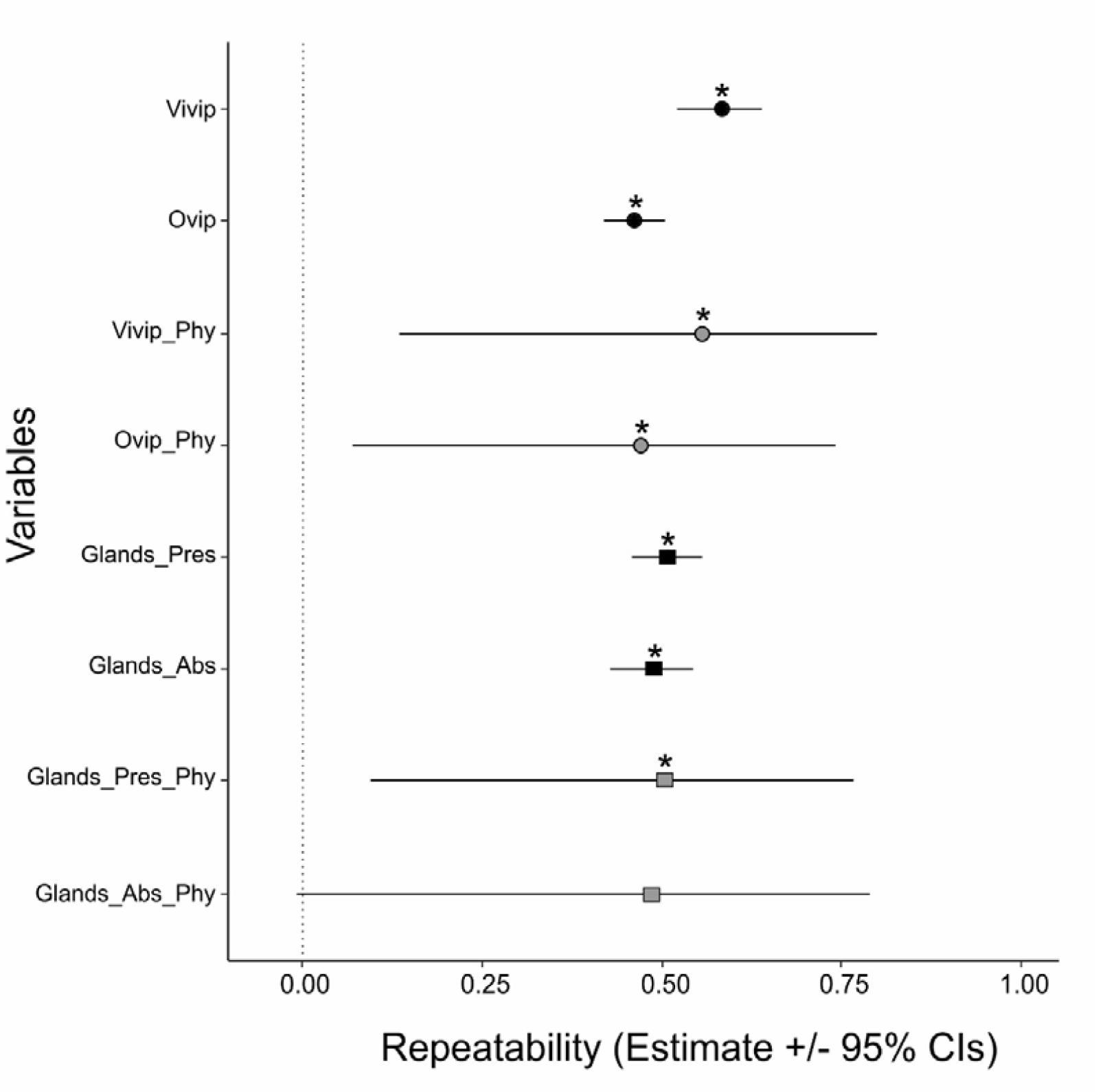
Repeatability values *R* (± 95% CI) for pooled data obtained for the different groups: occurrence of chemical glands groups (circles; absent= Glands_Abs; present= Glands_Pres) and reproductive mode (squares; viviparous= Vivip; oviparous = Ovip). The grey colour indicates that phylogeny (Phy) was corrected for the analysis, whereas the black colour indicates excluding the phylogenetic effect. The asterisk indicates statistically significance of the *R*-value.

**Table 4:**
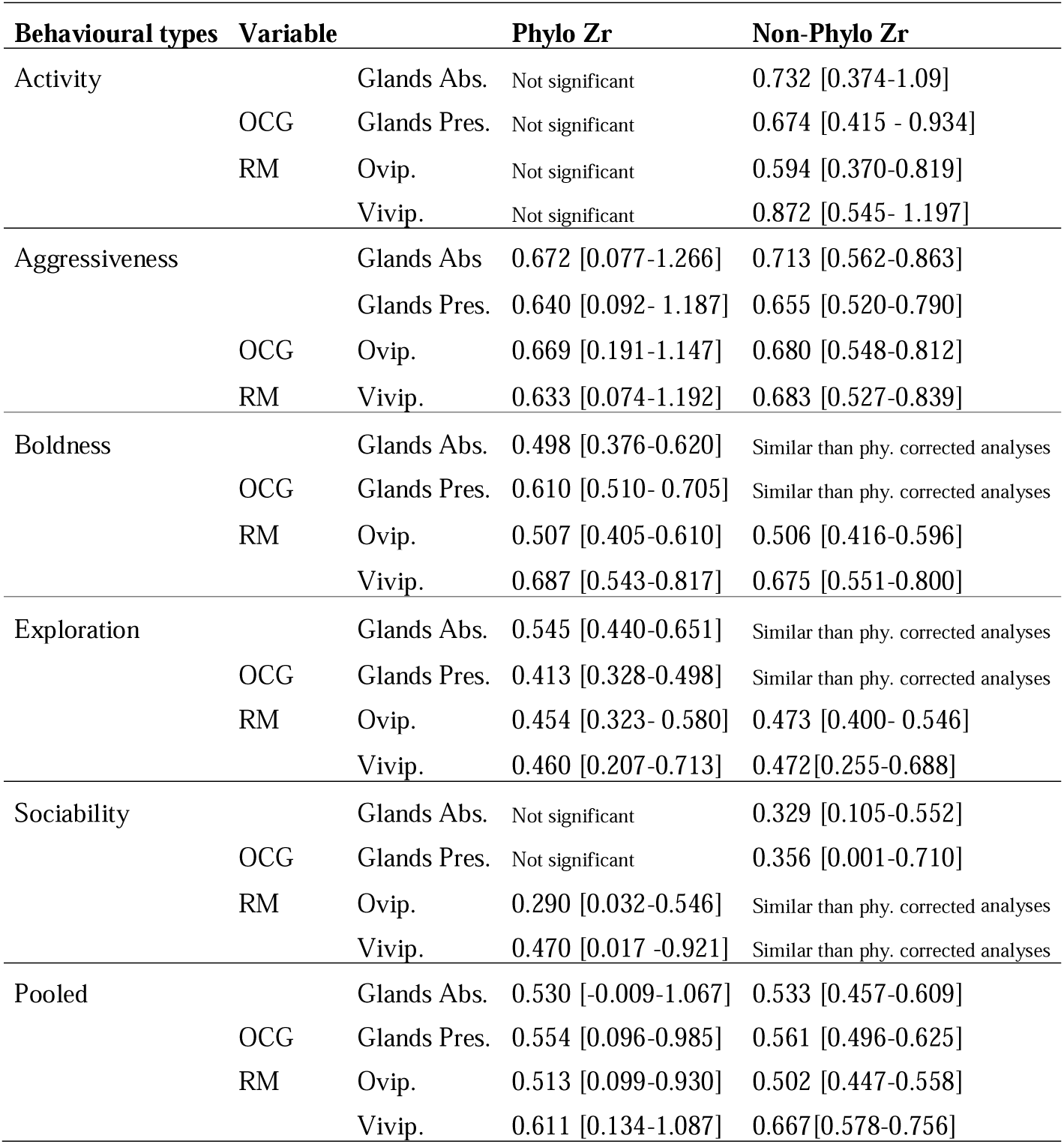
Effect size of repeatability values (Zr) and confidence intervals (95% CI) for the occurrence of glands (OCG: glands absent-glands present) and reproductive mode (oviparous-viviparous). Results for each behavioural type and for the pooled data, considering models with phylogenetic information (Phylo) and models without phylogenetic information (Non-Phylo).

**Table 5:** Repeatability values (*R*) and confidence intervals (95% CI) for the occurrence of glands (OCG: glands absent-glands present) and reproductive mode (oviparous-viviparous). Results for each behavioural type and for the pooled data, considering models with phylogenetic information (Phylo) and models without phylogenetic information (Non-Phylo).

| Behavioural types | Variables |  | *R* Phylo | *R* Non-Phylo |
| --- | --- | --- | --- | --- |
| Activity | OCG | Glands Abs. | Not significant | 0.624 [0.358-0.797] |
|  |  | Glands Pres. | Not significant | 0.588 [0.393-0.733] |
|  | RM | Ovip. | Not significant | 0.533 [0.354-0.675] |
|  |  | Vivip. | Not significant | 0.702 [0.497-0.833] |
| Aggressiveness | OCG | Glands Abs. | 0.586 [0.077-0.853] | 0.613 [0.510-0.698] |
|  |  | Glands Pres. | 0.565 [0.092-0.830] | 0.575 [0.478-0.659] |
|  | RM | Ovip. | 0.584 [0.188-0.817] | 0.592 [0.500-0.671] |
|  |  | Vivip. | 0.560 [0.074-0.831] | 0.594 [0.483-0.685] |
| Boldness | OCG | Glands Abs. | 0.461 [0.360-0.551] | Similar than phy. corrected analyses |
|  |  | Glands Pres. | 0.543 [0.470-0.608] | Similar than phy. corrected analyses |
|  | RM | Ovip. | 0.468 [0.384-0.544] | 0.468 [0.394-0.535] |
|  |  | Vivip. | 0.592 [0.495-0.674] | 0.592 [0.501-0.664] |
| Exploration | OCG | Glands Abs. | 0.497 [0.414-0.572] | Similar than phy. corrected analyses |
|  |  | Glands Pres. | 0.391 [0.317-0.461] | Similar than phy. corrected analyses |
|  | RM | Ovip. | 0.425 [0.317-0.523] | 0.441 [0.380-0.500] |
|  |  | Vivip. | 0.430 [0.204-0.613] | 0.440 [0.251-0.599] |
| Sociability | OCG | Glands Abs. | Not significant | 0.317 [0.104-0.502] |
|  |  | Glands Pres. | Not significant | 0.341 [0.001-0.611] |
|  | RM | Ovip. | 0.281 [0.032-0.498] | Similar than phy. corrected analyses |
|  |  | Vivip. | 0.438 [0.017-0.727] | Similar than phy. corrected analyses |
| Pooled | OCG | Glands Abs. | 0.485 [-0.008-0.788] | 0.488 [0.428-0.543] |
|  |  | Glands Pres. | 0.503 [0.096-0.766] | 0.508 [0.459-0.555] |
|  | RM | Ovip. | 0.472 [0.071-0.742] | 0.464 [0.420-0.506] |
|  |  | Vivip. | 0.544 [0.133-0.796] | 0.583 [0.521-0.639] |

The repeatability values for exploration showed significant, differences between both groups of OCG (Supplementary S2, Table 5). The repeatability for the “Glands Absent” group was higher (*R*_GAbs_= 0.497 [95% CI 0.414-0.572], Figure 5A, Table 5) than that of the “Glands Present” group (R_GPres_= 0.391 [95% CI 0.317-0.461], Figure 5A Table 5). Boldness behaviour evidenced an opposite pattern, with a higher repeatability for the “Glands Present” group (*R*_GPres_= 0.543 [95% CI 0.470-0.671], Figure 5A, Table 5) than for the “Glands Absent” group (R_GAbs_= 0.461 [95% CI 0.360-0.551], Figure 5 A, Table 5).

**Figure 5:**
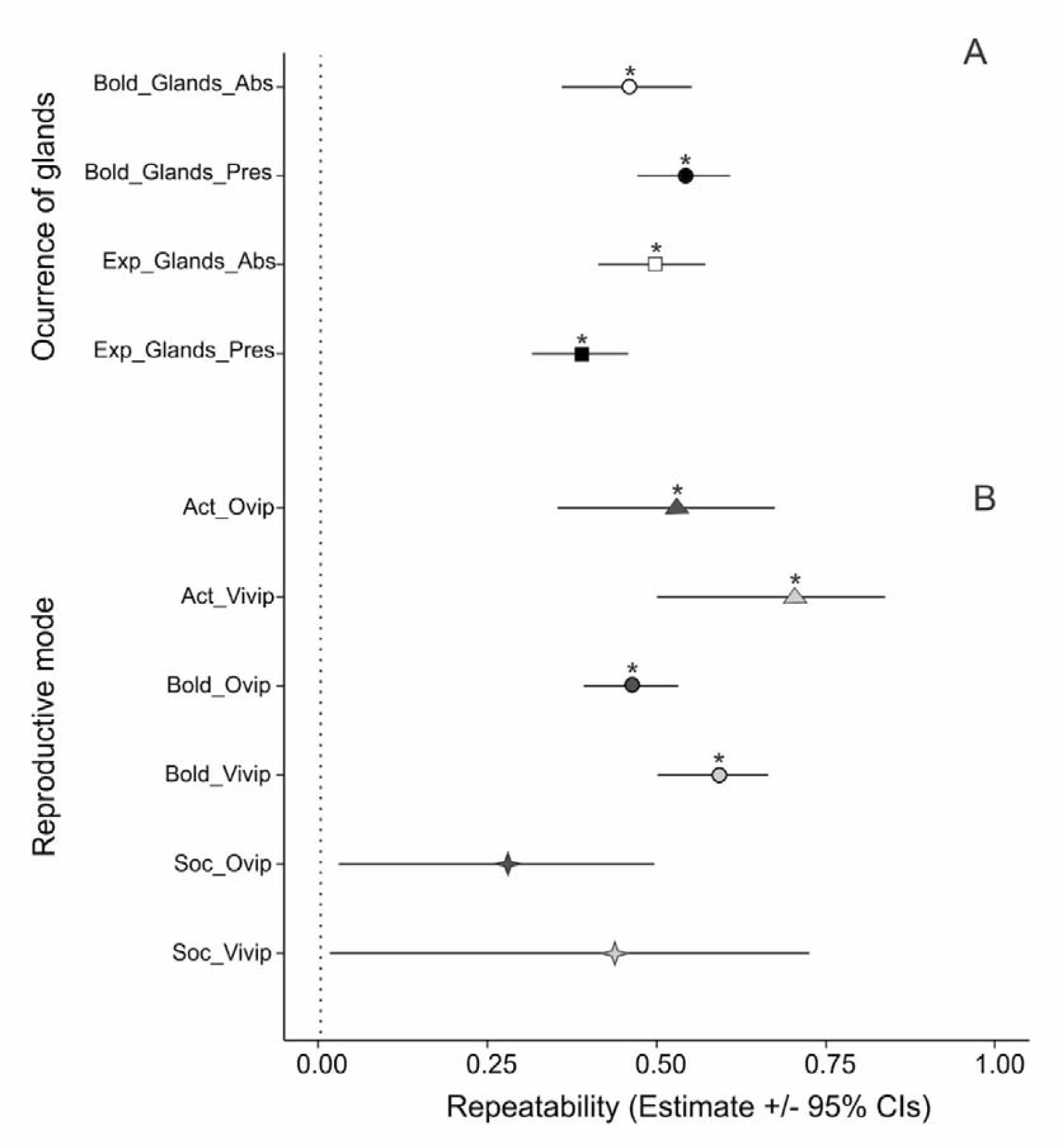
Repeatability values *R* (± 95% CI) for A-Occurrence of glands (black= absence, white= presence) and B-Reproductive mode (grey= oviparity, light grey= viviparity); considering activity (triangle), boldness (circle) and sociability (star). The asterisk indicates statistically significance of the *R-*value.

The slopes of the number of glands for phylogenetically corrected (*slope =* 0.0004 [95% CI −0.160-0.160] p = 0.99) and non-corrected analyses (*slope* = 0.013 [95% CI −0.036- 0.06] p = 0.60) were not significant.

The model considering reproductive mode was statistically significant (F _(df1= 2, df2= 35)_ = 3.530, p = 0.04). The σ^2^_total random_ was 0.299, with the phylogeny accounting for 77.36% of the random variability (σ^2^_phy_= 0.231), observations accounted for 19.80% of random variability (σ^2^_obs_ = 0.059) and fixed controlled variability explained 2.88% (σ^2^_fix. contr. var_ = 0.009). The *Zr* values for “Ovip” (p = 0.024; *Zr* = 0.513 [95% CI 0.099-0.930]; Table 4) and “Vivip” (p = 0.013; *Zr* = 0.611 [95% CI 0.134-1.087]; Table 4) were statistically significant. Posterior comparison did not detect any difference between the two groups (t = 0.63, p = 0.53). The overall repeatability for “Ovip” group was *R*_Ovip_= 0.472 [95% CI 0.071-0.742] (Figure 4; Table 5), while the repeatability for “Vivip” group was *R*_Vivip_= 0.544 [95% CI 0.133-0.796] (Figure 4; Table 5). The model without controlling for phylogenetic effects showed significant *Zr* for both the “Ovip” (p < 0.001; *Zr* = 0.502 [95% CI 0.447-0.558]; Table 4) and “Vivip” (p < 0.001; *Zr* = 0.667 [95% CI 0.578-0.756]; Table 4) groups. Posterior comparison detected differences between the two groups (t = 3.09, p < 0.01). The repeatability for “Ovip” group was *R*_Ovip_= 0.464 [95% CI 0.420-0.506] (Figure 4; Table 5), while for the “Vivip” group was *R*_Vivip_= 0.583 [95% CI 0.521-0.639] (Figure; Table 5).

The “Vivip” group showed higher repeatability values than for the “Ovip” group for activity, boldness, and sociability (Figure 5B, Table 5; Supplementary S2). Although the repeatability values for sociability behaviour were near zero, suggesting they were not statistically significant. Aggressiveness and exploration showed similar repeatability values for the two groups (Table 5).

## 4 DISCUSSION

Lizards exhibit varying degrees of sociability and animal personality, as was reported in different studies (e.g., Gardner et al., 2016; Waters et al., 2017). Meta-analytic studies are useful for integrating sociability indicators and animal personality data, allowing for the observation of patterns and trends across a broad phylogenetic context (e.g., Bell et al., 2009; Garamszegi et al., 2012). In this study, we evaluated the association of secretory glands and the reproductive mode of lizards with the repeatability of five behavioural types (activity, aggressiveness, boldness, exploration and sociability; details see Réale et al., 2007) obtained by compiling literature data. We found that the presence or absence of secretory glands did not affect repeatability values, but we observed higher repeatability values in viviparous than oviparous species. We did not find a correlation between the number of glands and repeatability data. In the following sections, we discuss the evolutionary implications of these results.

### 4.1 Influence of phylogeny on the repeatability of behaviour

Evolutionary relationships among species accounted for the greater variability in the pooled behavioural dataset, suggesting an important phylogenetic constraint on the repeatability of behaviour. Those closely related species show similar levels of individual behavioural consistency. In lizards, significant phylogenetic patterns were observed in life history (e.g., semelparity vs. iteroparity), reproductive strategies (K vs. r) and pace of life syndrome (slower vs. faster) (Dunham & Miles, 1985; Hallmann & Griebeler, 2015). Interestingly, these three traits were suggested as drivers of personality evolution (Réale & Dingemanse, 2010; Dammhahn et al., 2018; Laskowski et al., 2022). Strong phylogenetic influences were also reported for certain morphological (e.g., hemipenial morphology, Arnold, 1986; body size and number of scales, Oufiero et al., 2011; the number of secretory glands García-Roa et al., 2017) and behavioural traits (e.g., tongue flicks number, foraging and activity, Baeckens et al., 2017). Meta-analytic studies in other taxonomic groups have shown a strong influence of the phylogeny on animal personality in the analysed behaviours (e.g., Bell et al., 2009; Garamszegi et al., 2012). However, Dougherty and Guillette (2018), considering broader taxa sampling including insects, fish, reptiles, birds, and mammals, showed a shallow phylogenetic effect (10.71%) in the repeatability of five pooled behavioural types (i.e., activity, aggression boldness, exploration, and sociability). In more flexible behaviours, such as learning and cognition, the phylogenetic influence appears to be scarce (De Meester et al., 2022).

Phylogenetic constraints showed different patterns when the behavioural types were analysed separately. Phylogeny explained greater variation in the repeatability of activity and aggressiveness, whereas it explained less variation in the repeatability of boldness, exploration and sociability repeatability (Supplementary S2, Table 2). Activity refers to all movements made in both known and non-novel environments (Réale et al., 2007). While different taxonomic groups of lizards share similar activity patterns (Baeckens et al., 2017; but see Cooper Jr, 2005), there are groups considered more actives (e.g., Anguinomorpha, Lacertidae, Varanidae and Teiidae families) and groups less actives (e.g., Iguanidae, Geckkonidae and Scincomorpha;Vitt et al., 2003). Individual motivation for activity is also influenced by factors such as foraging mode, growth rate, physiological and metabolic processes, predator presence, seasonality, temperature, and age stage (e.g., Anderson & Karasov, 1981; Nicholson et al., 2005; Stamps, 2007; Lopez-Darias et al., 2012; Baeckens et al., 2017).

The animal personality framework quantifies aggressiveness by measuring aggressive displays and determining their repeatability (Réale et al., 2007; Laskowski et al., 2022). In lizards, aggressiveness is often studied in the context of male-male aggressions (e.g., Mouton & Wyk, 1993), and the displayed aggressive behaviours may exhibit phylogenetic conservatism (e.g., Jenssen, 1977; Kratochvíl & Frynta, 2002). Moreover, Dochtermann et al. (2015) demonstrated that aggressiveness has a stronger genetic basis than other behavioural responses, which could explain its strong phylogenetic pattern. Conversely, conspecific aggressive displays can be influenced by various factors, such as social status establishment, territorial disputes, and mating access (Carpenter, 1978; Fox et al., 2003), which may promote personality evolution (Réale et al., 2007; Laskowski et al., 2022). In crickets, individuals display higher aggressiveness and repeatability in the presence of conspecifics than in solitary conditions (Jäger et al., 2019). However, aggressiveness does not always promote individual consistency; Ruiz-Monachesi et al. (2023) showed that a more peaceful species, *Liolaemus albiceps*, exhibited behavioural consistency, whereas a more aggressive species, *L. coeruleus*, did not.

Boldness repeatability shows a weak influence of phylogeny, similar to other studies in lizards (e.g., Putman & Tippie, 2020) and in other groups (e.g., Garamszegi et al., 2012; Cauchoix et al., 2018). Boldness is strongly influenced by different fixed factors or moderators (details, see Tables 1 and 3), suggesting flexibility in this behaviour. De Meester et al. (2022), studied lacertid lizards for cognitive and learning processes and observed an absence of phylogenetic patterns and flexibility. Similarly, boldness appears to be a complex behaviour and responds to different factors in other taxa (e.g., Réale et al., 2007; Queller et al., 2022). Therefore, the weak phylogenetic constraint and the influence of various moderators suggest that boldness may evolve quickly (Lefebvre et al., 2016; Ducatez et al., 2017).

The absence of phylogenetic influence on sociability could be due to the limited representation of taxa in our dataset. We have data for only six species, four of which belong to the same family (Scincidae), while the other two belong to different distant families (Agamidae and Lacertidae; see Figure 1). Therefore, more studies with a broader species sampling are necessary to have a better understanding of the evolution of sociability personality in lizards. Furthermore, we found that aggressiveness and boldness exhibited the highest repeatability values, suggesting a possible counterbalance between phylogenetic constraints and flexible responses. This balance can interact with selective forces leading to the evolution of individual consistency in lizards.

### 4.2 The influence of secretory glands

Most lizard species are highly dependent on their chemosensory system (Mason & Parker, 2010) and possess chemical-emitting scents glands (exocrine glands), which secrete proteins and lipid-rich components (e.g., Mangiacotti et al., 2019).These glands are involved in various social interactions (Baird et al., 2015; Campos et al., 2017) and the number and diameter of these glands have been considered proxies for conspecific communication relevance (Martins et al., 2004; Baeckens, 2017; Ruiz-Monachesi et al., 2020). Baeckens and Whiting (2021) investigated the relationship between sociability and the evolution of chemical secretory glands and suggested that social grouping may drive investment in epidermal signalling glands in lizards. Sociability and conspecific interactions may also promote animal personality evolution (Laskowski et al., 2022), as suggested by the social niche specialization hypothesis (Bergmüller & Taborsky, 2010; Montiglio et al., 2013).

Contrary to our expectations, we did not find any association between the number of secretory glands and animal personality in lizards. However, when examining different behavioural types, we observed two contrasting patterns for exploration and boldness. Repeatability in exploration was higher in species without glands than in those species with glands. Exploration is defined as an individual’s tendency to move around in unfamiliar environments (i.e., neophilia; Réale et al., 2007). This pattern could be explained by the high-investment signal hypothesis, where a poorly developed or absent chemical emissary system would be counterbalanced by an efficient receiving system and higher exploration behaviour (Ruiz-Monachesi et al., 2020, 2022). For instance, different *Liolaemus* species with few or no precloacal pores showed higher exploration behaviour than species with a greater number of precloacal pores (Ruiz-Monachesi et al., 2020). Therefore, if the secretory glands are absent, individuals may perceive their environment as more novel when exploring, potentially contributing to the development of individual consistency in this behaviour. Furthermore, it is important to note that the secretions of femoral/precloacal glands convey identity-related information in lizards (e.g., Cooper et al., 1999; Mangiacotti et al., 2019). The depositions of these scents may decrease an individual’s perception of novelty in a new environment and consequently impact exploration behaviour.

We observed an opposite pattern for boldness, higher repeatability in species with chemical emissary glands than those without glands. Boldness can be defined as an individual’s tendency to take risks, measured across different contexts such as basking, foraging, novelty-exposure, predators’ attacks, and social contexts (Réale et al., 2007). A possible explanation for our results could be due to the influence of the endocrine system on animal personality. Intrinsic factors, such as hormones, present associations with animal personality and individuals that are relatively bolder may also present relatively high hormone levels (Niemelä & Dingemanse, 2018). Testosterone has been shown to increase an individual’s boldness behaviour (e.g., Raynaud & Schradin, 2014) and there is an association between boldness behaviour and testosterone receptors presence (Kabelik et al., 2022). In lizards, testosterone can activate the secretion of femoral glands in adults of both sexes (Hews & Quinn, 2003). In fact, femoral glands’ activity is mediated by several androgens in Squamata (e.g., Abell, 1998; Hews & Quinn, 2003). Therefore, the integration of hormonal influence, behaviour, and morphology may explain the observed pattern.

### 4.3 The influence of viviparity

In lizards, viviparity has been associated with hard climatic conditions such as high altitudes, hypoxia, and cold climates (Watson & Cox, 2021; Zimin et al., 2022). Generally, embryos in a maternal environment are exposed to more stable conditions than those developing in eggs (e.g., Deeming, 2004; Cruz et al., 2022). However, some maternal behaviour, such as basking opportunity, affect juvenile phenotype in viviparous lizards (Wapstra, 2000). Early life experiences, such as those that occur during embryonic development, influence an individual’s personality (e.g., Siviter et al., 2017a). Therefore, viviparous and oviparous species experience development differences that could favour distinctions in the degree of animal personality between them, suggesting that viviparity promotes the evolution of animal personality. Two possible explanations for this difference could be mentioned. First, different social characteristics may be associated with viviparity, which appears to promote the evolution of sociability in these taxa (Gardner et al., 2016; Halliwell et al., 2017). In fact, all those early social experiences experimented by individuals after birth may favour individual behavioural consistency in some viviparous family-living lizard species (Riley et al., 2017). Second, temperature variations during egg incubation impact the phenotype (behaviour, morphology, physiology, sex) of oviparous reptiles (Deeming, 2004; Noble et al., 2018). Some flexible behaviours, such as cognition in a social context, are affected by differences in incubation temperatures; individuals raised in colder temperatures learn a social task more quickly or are more active than those raised in warmer temperatures (Dayananda & Webb, 2017; Siviter et al., 2017b; but see Amiel & Shine, 2012). The temperature of incubation also affects boldness in oviparous lizard species; individuals raised in warmer temperatures are bolder than those from cooler temperatures (Siviter et al., 2017a). In oviparous skinks lizards, warmer incubation temperatures are associated with higher personality consistency in different behavioural traits (activity, exploration, sociability, and boldness) than those incubated at lower temperatures (de Jong et al., 2022). Natal temperature differences can also explain individual differences in dispersal (exploration) behaviour in lizards (Bestion et al., 2015).

### 4.4 Publication bias

We observed asymmetry in the funnel plot, which may suggest the presence of publication bias in the dataset, however, this asymmetry and publication bias are not always associated (Sterne et al., 2005). A funnel plot indicates the precision in the estimation of the underlying treatment effect, and it increases as the sample size of the studies in the review increases. As was summarized by Sterne et al. (2005), asymmetry may come from different sources as: i-selection biases (due to location, language, citation, etc.); ii-true heterogeneity (the size of the effect differs according to study size); iii-data irregularities (poor methodological design of small studies, inadequate analysis, fraud); iv-artefact (heterogeneity due to poor choice of effect measure) or v-simple chance. The asymmetry observed here, may be due to a true heterogeneity, as the compiled data had a wide range of sample sizes (min= 3, max= 807). Another possibility for the observed asymmetry could be due to an artefact associated with the nature of the data. For reptiles, was showed a mean repeatability around 0.50, while for other taxa, such as mammals, repeatability would be higher (0.65-0.70; Bell et al., 2009). This suggests that for lizards, higher *R* values than 0.50 are less frequent and the potential publication bias could be an effect associated with the low frequency of research reporting higher than 0.50 or 0.60 repeatability values. Another possible explanation for the low frequency of higher than 0.50 repeatability values could be associated with the observed taxonomic bias. The majority of studies were made in a few species as those species belonging to Scincomorpha (e.g., *Lampropholis delicata*), followed by Lacertoidea and Iguania. Others, such Gekkota and Platynota were represented with one or two studies. This taxonomic bias would favour decreasing the value of repeatability effect size. An increase in the taxonomic bias could favour an increase in the between individuals’ variability, decreasing the mean of effect size repeatability. Even if we considered interspecific variations, within-population variations also contribute to the low repeatability of data (Ives et al., 2007).

We do not discard the possibility of having missed studies due to a sampling bias, however, different parameters obtained here (e.g., *Q*, *τ*^2^, *I*^2^, *H*^2^) suggest more variability present than those expected by a sampling bias (Rothstein et al., 2005). This was supported by the high value of Rosenthal’s failsafe (1,114,233), the results of the meta-analysis can be considered robust to publication bias because it would be very unlikely to find that additional number of unpublished results (Becker, 2005).

### 4.5 Conclusions

Nowadays, there were described ∼ 7,310 lizard species in the world (Uetz et al., 2023), of which we compiled personality data of 37 species. These species represent 0.50% of lizard diversity. Some behaviours such as sociability, activity and aggressiveness have been less studied, thus our study evidenced the presence of taxonomic bias considering animal personality in lizards. In spite of this asymmetry, phylogeny and studies-observations explained a significant proportion of the random heterogeneity in lizard’s personality. Our study suggests that animal personality traits such as activity and aggressiveness evolve slowly due to phylogenetic constraints. However, traits such as boldness, exploration, and sociability may have faster evolutionary rates, as they showed weak phylogenetic influence.

The absence of chemical emissary glands appears to be associated with greater repeatability in exploration. Finally, viviparity reproductive mode could be associated with the animal personality in lizards, possibly due to the existence of more stable internal conditions during development and/or to posterior early-life experiences, especially in highly sociable species. This study represents a first step to understanding the interplay among the lizards’ animal personality, life-history traits (reproductive mode) and morphology (chemical emissary glands) under a broad evolutionary context.

### Supplementary data

https://data.mendeley.com/datasets/3zgzkddrjd/1

## Author Contributions

**Mario R. Ruiz-Monachesi:** Conceptualization (equal), data curation (lead), formal analysis (lead), investigation (equal), methodology (equal), writing-original draft (equal), writing-review and editing (equal), funding acquisition (equal), supervision (supporting), project administration (supporting). **Juan J. Martínez:** Conceptualization (equal), data curation (supporting), formal analysis (supporting), investigation (equal), methodology (equal), writing-original draft (equal), writing-review and editing (equal), funding acquisition (equal), supervision (lead), project administration (lead).

## Acknowledgements

We thank J.J. Wiens and J.J. Cuervo, who provides the Squamate molecular phylogeny and secretory pores data for *Intellagama lesueurii*, respectively. The authors are career researchers of the Consejo Nacional de Investigaciones Científicas y Técnicas (CONICET). This study was partially funded by CONICET (PIP 2022-2024) to JJM, MRR-M, and Lucía Sommaro.

## All authors have seen and approved the manuscript

### Competing Interests

The authors declare there are no competing interests.

### Data Availability

Tables and figures were uploaded in the context of this manuscript. Data matrix and scripts will be uploaded to a repository under acceptance of the manuscript.

## Notes

### Competing Interest Statement

The authors have declared no competing interest.

